# Genetic architecture of a light-temperature coincidence detector

**DOI:** 10.1101/2024.10.30.621125

**Authors:** Adam Seluzicki, Joanne Chory

## Abstract

Light and temperature variations are inescapable in nature. These signals provide daily and seasonal information, guiding life history determinations across many taxa. Here we show that signals from the PHOTOTROPIN2 (PHOT2) blue photoreceptor combine with low temperature information to control flowering. Plants lacking PHOT2 flower later than controls when grown in low ambient temperature. This phenotype is blocked by removal of NON-PHOTOTROPIC HYPOCOTYL 3 (NPH3) and recapitulated by reducing blue light intensity or removing the transcription factor CAMTA2. PHOT2 and CAMTA2 show non-additive genetic interactions in phenotype and gene expression. Network-based co-expression analysis indicates system-level control of key growth modules by PHOT2 and CAMTA2. CAMTA2 is required for low temperature up-regulation of *EHB1*, a known NPH3-interacting protein, providing a mechanism of temperature information input to the PHOT-NPH3 blue light signaling system. Together these data describe the genetic architecture of environmental signal integration in this blue light-low temperature coincidence detection module.

## INTRODUCTION

Daily and seasonal cycles of light and temperature provide information that organisms transform into modulation signals, tuning physiology and development to the environment^1–7^. Light quality, intensity, duration, and direction are sensed in parallel by a suite of photoreceptor proteins^8^. Temperature sensing involves many systems including calcium currents, membrane fluidity, RNA secondary structure, histone dynamics, and the photoreceptors themselves^9–15^. Light and temperature are processed by plants simultaneously, with the combination of light and temperature conditions providing more information than either stimulus alone. Physiological and developmental functions that demonstrate differential sensitivity to particular combinations of environmental light and temperature are of interest to understand mechanisms of processing environmental information.

PHOTOTROPINs (PHOTs) are blue-light activated kinases, localized to the plasma membrane in complexes with accessory signaling factors^16^. PHOT1 is more sensitive than PHOT2^14,17^. Both participate in phototropic bending, chloroplast movement, leaf flattening, and stomatal opening^18–21^. Several members of the NON-PHOTOTROPIC HYPOCOTYL 3 / ROOT PHOTOTROPISM 2 (NPH3 / RPT2)-Like (NRL) family of BROAD-TRAMTRACK-BRICK-A-BRACK (BTB)-domain signaling proteins participate in PHOT-dependent processes. NPH3 and RPT2 contribute to phototropic bending, RPT2 and NCH1 function in chloroplast movement, and all three are involved in controlling leaf curvature^22^. While the other photoreceptor families have well-described links to regulation of flowering, there have been few indications that PHOT signaling components contribute to this developmental transition^18^.

<u>CA</u>L<u>M</u>ODULIN-BINDING <u>T</u>RANSCRIPTIONAL <u>A</u>CTIVATOR (CAMTA) transcription factors are wide-spread in eukaryotes. They are involved in neural and cardiac development and signaling in animals, and regulate immunity, drought, salt, and hormone responses in plants^23–28^. In *Arabidopsis,* CAMTA3 is the best studied due to its role in regulating immunity in response to low temperature, acting redundantly with CAMTA1 and 2^24,25,29^. There are conflicting reports regarding a possible interaction between CAMTA3 and the NRL protein NCH1^26,30^. Temperature regulation of CAMTA3 function has been mapped to the DNA binding domain^31^. Other CAMTAs have not been studied in as much detail. Outside of calcium, this family of transcription factors has not been directly linked to the regulation of known signaling pathways in plants.

Here we examine the PHOTOTROPIN contribution to flowering in the context of environmental light-dark cycles and ambient temperature. We show that NPH3 functions at the center of a coincidence detector, integrating blue light intensity from PHOT2 and low ambient temperature from CAMTA2. We provide genetic and molecular evidence that PHOT2 and CAMTA2 work together to fine-tune flowering and regulate highly overlapping gene sets. Network co-expression analysis shows system-level mis-regulation of fundamental growth control modules in *phot2* and *camta2* mutants. Low temperature- and CAMTA2-dependent expression of the NPH3-interacting protein EHB1 is a likely point of temperature information input to the PHOT-NPH3 light signaling pathway. Thus, PHOT2, CAMTA2, and NPH3 function together to integrate light and temperature information, buffering development against environmental conditions.

## RESULTS

### PHOT2 and NPH3 control coordinated light and temperature sensitivity

We analyzed the relationship between PHOTOTROPIN signaling and flowering under 16h light:8h dark (LD16:8, light intensity: ∼100μmol/m^2^s) at constant temperature, either 20°C or 15°C [see spectrum in Extended Data 1A]. We found that two independent alleles of *PHOT2* (*phot2* = *phot2-101* (SALK_142275), and *phot2 gl1* = *phot2-1 gl1*) showed a consistent increase in the number of leaves at flowering (Total Leaf Number, TLN) relative to controls [Fig. 1A-B]. While we observed a slight difference at 20°C, this effect was particularly pronounced at 15°C. Two alleles of *phot1* in these respective genetic backgrounds did not show consistent differences in leaf number relative to controls [Fig. 1A-B]. We tested other known PHOT signaling components. We found that *nph3* showed slightly reduced TLN relative to control [Fig. 1A]. Two mutants with strong effects on chloroplast movement, *chup1* and *jac1*, were similar to controls, indicating that chloroplast movement is unlikely to be responsible for the phenotype [Fig. 1A].

**FIGURE 1:**
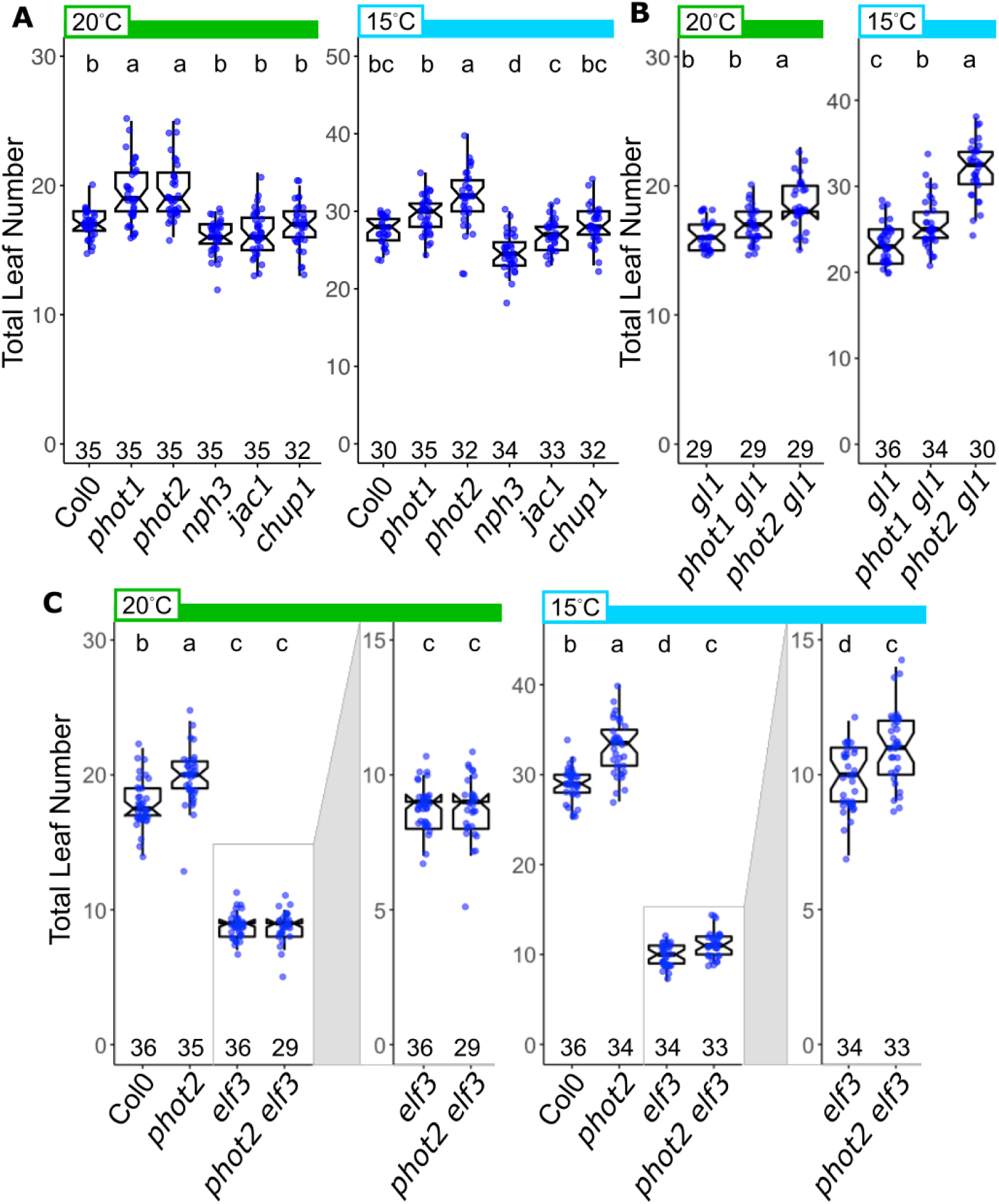
PHOTOTROPIN signaling regulates temperature-labile flowering. (A-C) Flowering assayed by total leaf number at the time of bud appearance in the indicated genotypes grown in the indicated temperature conditions (20°C - green bars, 15°C - light blue bars). Light conditions: 16 light:8h dark (LD16:8) photoperiod, light intensity: ∼100-110 μmol m-2 s-1 (see supplemental Fig 1). Box plots show the median, 25th and 75th percentiles, and whiskers extending to 1.5*interquartile range (IQR). Notches approximate the 95% confidence interval of the median. N for each genotype/condition is indicated along the x-axis. Different letters indicate statistically significant difference between groups (α=0.05), within each temperature condition by one-way ANOVA+Tukey HSD. N for each genotype is noted just above the x-axis. (A) Total leaf number at flowering in PHOT-related signaling factors. (B) Total leaf number at flowering in *phot1* and *phot2* mutants in the Col-0 *gl1* mutant background. (C) Total leaf number at flowering in Col-0, *phot2*, *elf3*, and *phot2 elf3*. Y axes for *elf3* and *phot2 elf3* boxplots are expanded for each temperature in boxes to the right of each panel for clarity.

The circadian evening complex component ELF3 is a central player in the floral transition as a repressor of the florigen FT, and is a temperature sensor itself, with the mutant showing reduced sensitivity to cool ambient temperature^32,33^. We therefore asked if the hypersensitivity to low temperature in *phot2* would persist in the absence of ELF3. We found that *elf3* and *phot2 elf3* were identical at 20°C, both flowering with fewer leaves than Col-0 and *phot2* [Fig. 1C]. However, at 15°C, *phot2 elf3* flowered with more leaves than *elf3*. Thus, temperature sensitivity unmasked in the *phot2* mutant occurs downstream of, or in parallel with, ELF3 function.

We then tested an expanded set of temperature and light conditions. We examined *phot2* mutants under 25°C, finding that the mutants were indistinguishable from Col-0 controls, while again observing increased TLN in *phot2* under 15°C [Fig. 2A, white panels]. We filtered out the blue part of the spectrum (Yellow Filter, YF) and found that Col-0 plants behave identically to *phot2* under 15°C [Fig. 2A, yellow panels, Extended Data 1B]. Calculating the rate of leaf generation as Leaves per Day (LPD), we found that both the number of leaves and the rate of leaf generation were identical in Col-0 and *phot2* plants in 15°C YF vs white light (WL) [Fig. 2B]. Leaf Number Ratio (LNR), the ratio between TLN in 15°C and 25°C, was larger in *phot2* than Col-0 in WL, but similar between the genotypes in YF, matching *phot2* in WL [Fig. 2C]^34^. Increasing the blue or red components of low WL (∼50μmol/m^2^s) did not modify the high TLN phenotype of *phot2* plants in 15°C, suggesting that the phenotype derives from the intensity of blue light up to a point of saturation, rather than blue:red ratio or other relative proportions in the spectrum [Extended Data 1C-D, 2A-B].

**FIGURE 2:**
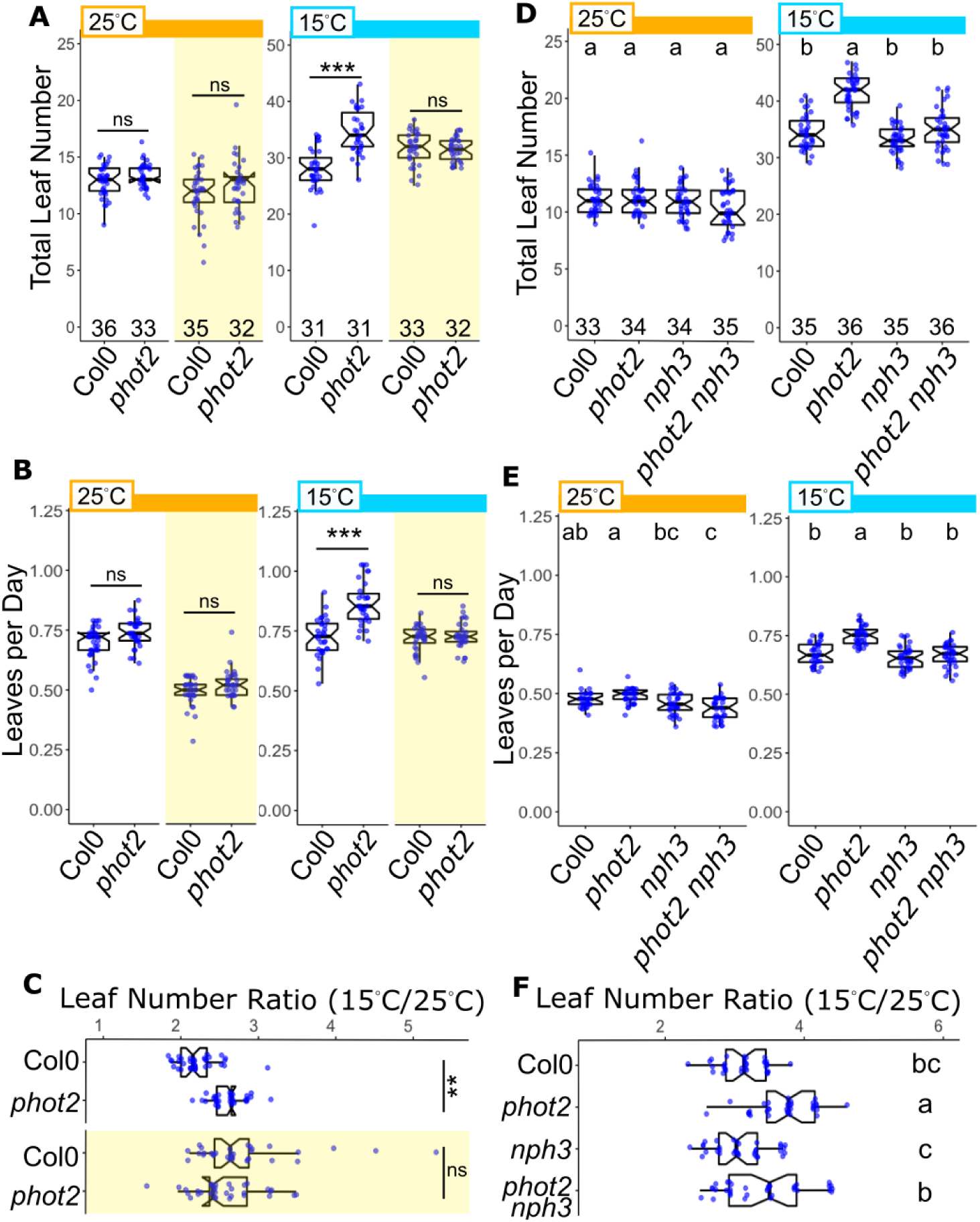
PHOTOTROPIN2 integrates blue light and temperature signals in a light intensity- and NPH3-dependent manner. (A,C) Total leaf number at flowering under indicated light and temperature conditions (25°C - orange bars, 15°C - light blue bars). See Extended Data 1 for representative spectra. Data are presented as in Fig. 1. (A) Flowering assayed under white light (WL, ∼100-110 μmol m^-2^ s^-1^ - white background) + yellow filtered light (YL - yellow shaded regions), which specifically reduces the blue component. (B) Leaves per Day (Total Leaf Number / Days Post-Germination) calculated from the experiments in (A). (C) Total leaf number in Col-0 control, *phot2*, *nph3*, and *phot2 nph3* mutants under low white light (∼50-55 μmol m^-2^ s^-1^). (D) Leaves per Day calculated from the experiments in (B) (E,F) Leaf number ratio temperature responses calculated from (A) and (C), respectively. (A,B,E) Two-way ANOVA (genotype X light) result is shown. ns = not significant at p=0.05, *** p<0.001, ** p<0.01

Given that PHOT2 is known to work with NPH3, and that *phot2* and *nph3* showed opposing TLN phenotypes [Fig. 1A], we asked if *nph3* might be epistatic to *phot2*. We examined Col-0, *phot2*, *nph3* and *phot2 nph3* under 25°C and 15°C and assayed TLN at flowering (pooled WL data from the experiments in Extended Data 2A-B). All four genotypes were indistinguishable under 25°C. Under 15°C, *phot2* showed expected increased TLN, *nph3* was similar to Col-0, and *phot2 nph3* showed clear suppression of the *phot2* phenotype [Fig. 2D]. Interestingly, *nph3* showed its own differential light sensitivity under 15°C; *nph3* flowered with a similar TLN as Col-0 under low WL (50μmol/m^2^s), but with increasing light intensity (100μmol/m^2^s) it showed reduced TLN relative to control [Extended Data 2C]. This suggests that NPH3 acts to counter light intensity-dependent promotion of flowering. Together, these data strongly support the hypothesis that PHOT2 acts with NPH3 as part of a light-temperature signaling integration system, using the combination of blue light intensity and low temperature information to buffer or fine-tune development, likely by antagonizing parallel light signals that feed into flowering time control pathways.

### Non-additive genetic interaction between PHOT2 and CAMTA2

We performed an RNA-sequencing experiment using *gl1*, *phot1 gl1*, and *phot2 gl1*, grown under either 20°C or 15°C and sampled at lights-off (ZT16) and the following lights-on (ZT24) [Supplementary Data 1]. We identified differentially expressed genes (DEGs), examined the promoters of these genes using ELEMENT, finding that the promoters of DEGs in *phot2* in 15C at ZT16 were significantly enriched for variants of CAMTA transcription factor binding motifs (CGCG) [Extended Data 3A-B]^35,36^. Examination of available DNA Affinity Purification (DAP)-seq data revealed enrichment for CAMTA1 and CAMTA5 binding events among these genes [Extended Data 3C]^37^. None of the 75 other TFs with a similar number of reads to CAMTA1 in this database showed similar enrichment, indicating a highly specific effect [Extended Data 3D].

We screened a panel of *camta* mutants in our flowering assay. We found increased TLN in lines with the *camta2* mutation, with the strongest effect in the *camta1 camta2* double mutant [Fig. 3A]. Mutations in *camta1* and *camta3* did not correlate with increased TLN. Our data thus implicate *CAMTA2*, with possible redundant contribution of *CAMTA1*, as the primary members of this group regulating TLN at flowering. We hypothesized that *PHOT2* and *CAMTA* genes may interact genetically. We observed that *phot2 camta1* plants flower with more leaves than *camta1* alone, an effect similar to *phot2* vs Col-0 control [Fig. 3B]. However, *phot2* failed to increase leaf number in the *camta2* background, with *phot2*, *camta2*, and *phot2 camta2* being statistically indistinguishable under either growth temperature. We also found *phot2 camta1 camta2* TLN to be the same as *camta1 camta2*. We asked if the *camta2* mutant would respond to warmer temperature in a manner similar to *phot2*. We assayed flowering at 25°C and 15°C, observing that all genotypes grown under 25°C show indistinguishable TLN at flowering, but *phot2*, *camta2*, and *phot2 camta2* show more leaves than control under 15°C [Fig. 3B, Extended Data 4]. Together, these data demonstrate that PHOT2 and CAMTA2 regulating temperature sensitivity of flowering. The non-additive genetic interaction between PHOT2 and CAMTA2 suggests that these factors operate in a coordinated manner.

**FIGURE 3:**
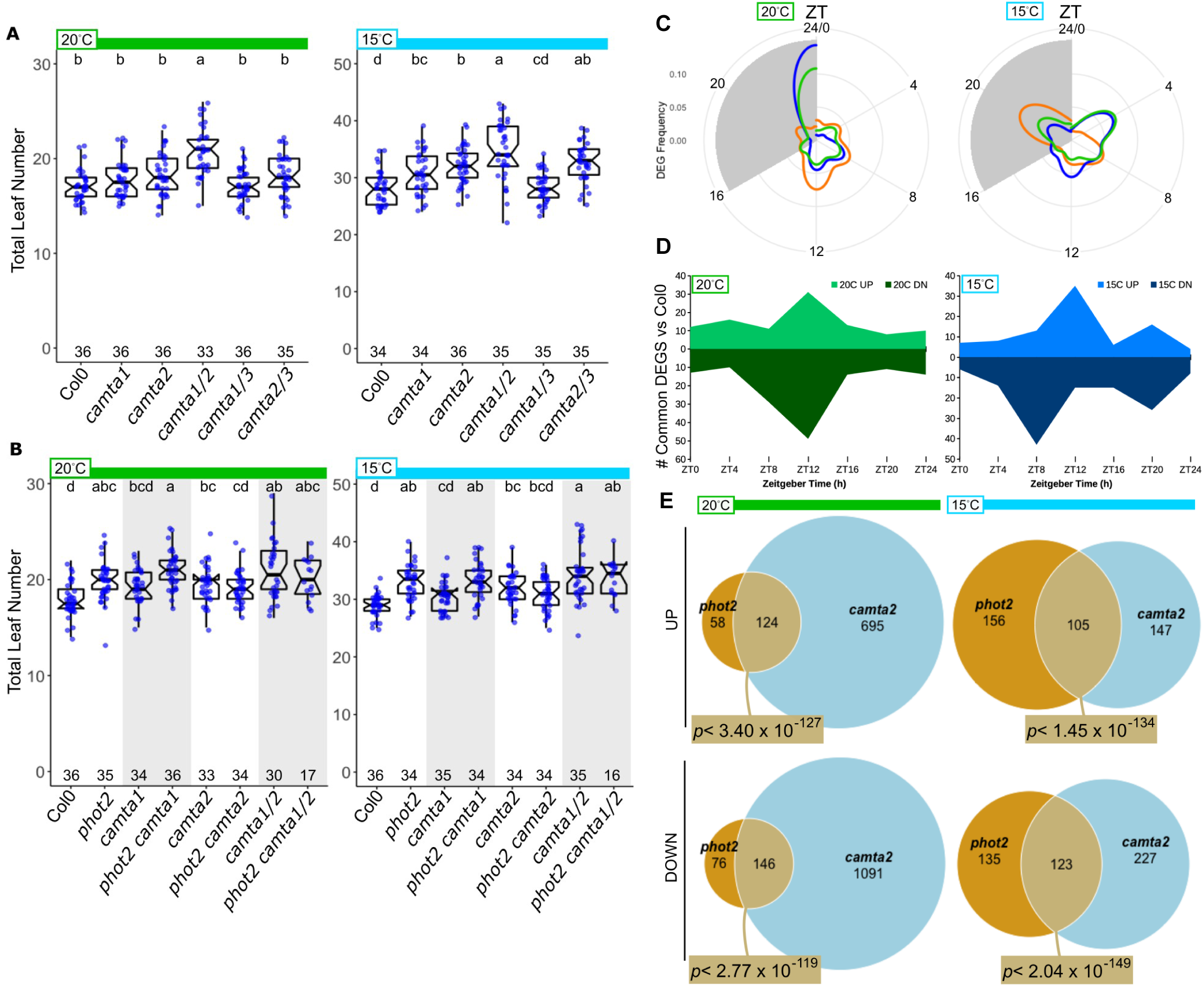
Non-additive genetic interaction between *phot2* and *camta2*. (A,B) Total leaf number at flowering under indicated temperature conditions (20°C - green bars, 15°C - light blue bars), as in Figure 1. The *camta2-1* allele is used in A-E. (A) Flowering assayed in *camta1, camta2, and camta3* single and multiple mutants. (B) Flowering assayed in *camta1* and *camta2* mutants +/- *phot2.* Pairs of genotypes in white and shaded areas indicate key comparisons. (C-E) Col-0, *phot2*, *camta2*, and *phot2 camta2* plants were grown on soil at either 20C or 15C. Samples were collected every four hours, starting at ZT0 (lights-on) of day 10 for RNA sequencing. Differentially expressed (DE) genes were identified between genotypes, within each timepoint and temperature. Threshold for DE was |0.25| fold-change, FDR< 0.05. (C) The number of genes identified as differentially expressed in *phot2*, *camta2*, and *phot2 camta2* plants compared to Col-0 control within each timepoint and temperature, expressed as a circular histogram to emphasize time of day. The x-axis (clockwise around the circle) is in Zeitgeber Time (ZT) where ZT0 is the time of lights-on, ZT16 is lights-off, and ZT24 is the next lights-on. The grey region indicates the dark period (ZT16-24). (D) Graphs show the number of up- or down-regulated DE genes identified in common in *phot2*, *camta2*, and *phot2 camta2* plants, plotted by time point and temperature. Lighter color (above 0) indicates up-regulated genes, and the darker color (below 0) indicates down-regulated genes at each time point across the day. Some DE genes were identified as differentially expressed at multiple time points, and are therefore counted in more than one timepoint (see Supplementary Data). (E) Venn diagrams of UP- and DOWN-regulated genes identified in *phot2* and *camta2* mutant plants relative to Col-0. DE gene lists for each genotype across the day were collapsed into a single list, and compared between genotypes. P-values indicate the significance of overlap by Fisher’s exact test (19,139 total genes after expression level filtering).

### PHOT2 and CAMTA2 co-regulate gene expression across the day

We carried out a full day time course RNA-seq analysis, sampling Col-0, *phot2*, *camta2*, and *phot2 camta2* plants every four hours. To ensure close molecular correlates with our flowering assay, we sampled aerial tissue from soil-grown plants^38^. *PHOT2* and *CAMTA2* expression patterns across the day matched the Col-0 control in *camta2* and *phot2* mutants, respectively [Extended Data 5A], arguing against the hypothesis of transcriptional feedback between these genes.

The temporal occurrence of DEGs identified in *phot2, camta2*, and *phot2 camta2* were similar, with a high proportion identified in the mid-day [Fig. 3C]. Examining the DEGs identified in common among all three mutant lines across the day (common DEGs), we observed that genes that were misregulated tended to be identified particularly in the mid-day under both 20°C and 15°C, with down-regulation under 15°C more often observed sightly earlier than up-regulation (ZT8 vs ZT12) [Fig. 3D]. DEGs identified in *phot2*, *camta2*, and *phot2 camta2* plants were highly overlapping within each time point, with evidence of more misregulated genes in the double mutant [Extended Data 6]. We found highly significant overlap of genes regulated in the same direction (up or down relative to Col-0 control) in *phot2* and *camta2* in both temperatures [Fig. 3E]. We cannot make a definitive claim as to which genes are directly regulated by CAMTA2. However, examination of the promoters of the genes identified as common DEGs in *phot2*, *camta2*, and *phot2 camta2* under 15°C using the Motif Finder tool of ELEMENT, 43 genes contain the consensus CAMTA-binding motif (CGCG) and 126 genes contain the CAMTA2 variant motif CGTG, often in multiple instances within 1kb of the transcription start site [Extended Data 5B, Supplementary Data 2]^35,36,39^.

We reasoned that system-level regulation may be more visible when analyzed across the complete time series, rather than single time-point comparisons. We identified co-expressed “communities” of genes in our data set [Fig. 4A, Supplementary Data 3, Methods]. Mean scaled expression levels of each gene in the resulting communities were averaged and communities 3, 6, and 15 are shown [Fig. 4B]. We compared the scaled expression in *phot2*, *camta2*, and *phot2 camta2* to Col-0 at each time point to identify communities that were systematically mis-regulated, finding time- and temperature-dependent differences [Fig. 4C]. Gene Ontology analysis of these communities reveals system-level regulation of specific processes by PHOT2 and CAMTA2, and provide clues as to potential pathways that may control TLN. Community 3 shows global up-regulation of ribosome biogenesis-related processes [Fig. 4D]. This pattern persists when limiting the gene set to the GO category Ribosome Biogenesis [Fig. 4E]. This community is more strongly up-regulated in *camta2* and *phot2 camta2* than in *phot2*. Community 6 shows up-regulation of cell cycle-related processes across the day in the mutants, and the pattern is maintained limiting the gene set to GO: Cell Cycle [Fig. 4F,G]. Community 15 shows up-regulation of DNA replication related processes in the mutants, most clearly in the evening and night time points [Fig. 4H]. While there is only one statistically different time-point seen in Community 15 at 15°C as identified by the network [Fig. 4C], refining it to the members of the GO category DNA Replication reveals clear up-regulation in the mutants [Fig. 4I]. These trends are preserved in the Ribosome Biogenesis, Cell Cycle, and DNA Replication groups when examined at the single gene level [Extended Data 7]. This unbiased network building and analysis strategy allows us to resolve system-level effects on fundamental growth-related modules by PHOT2 and CAMTA2 [Fig. 4E, G, I].

**FIGURE 4:**
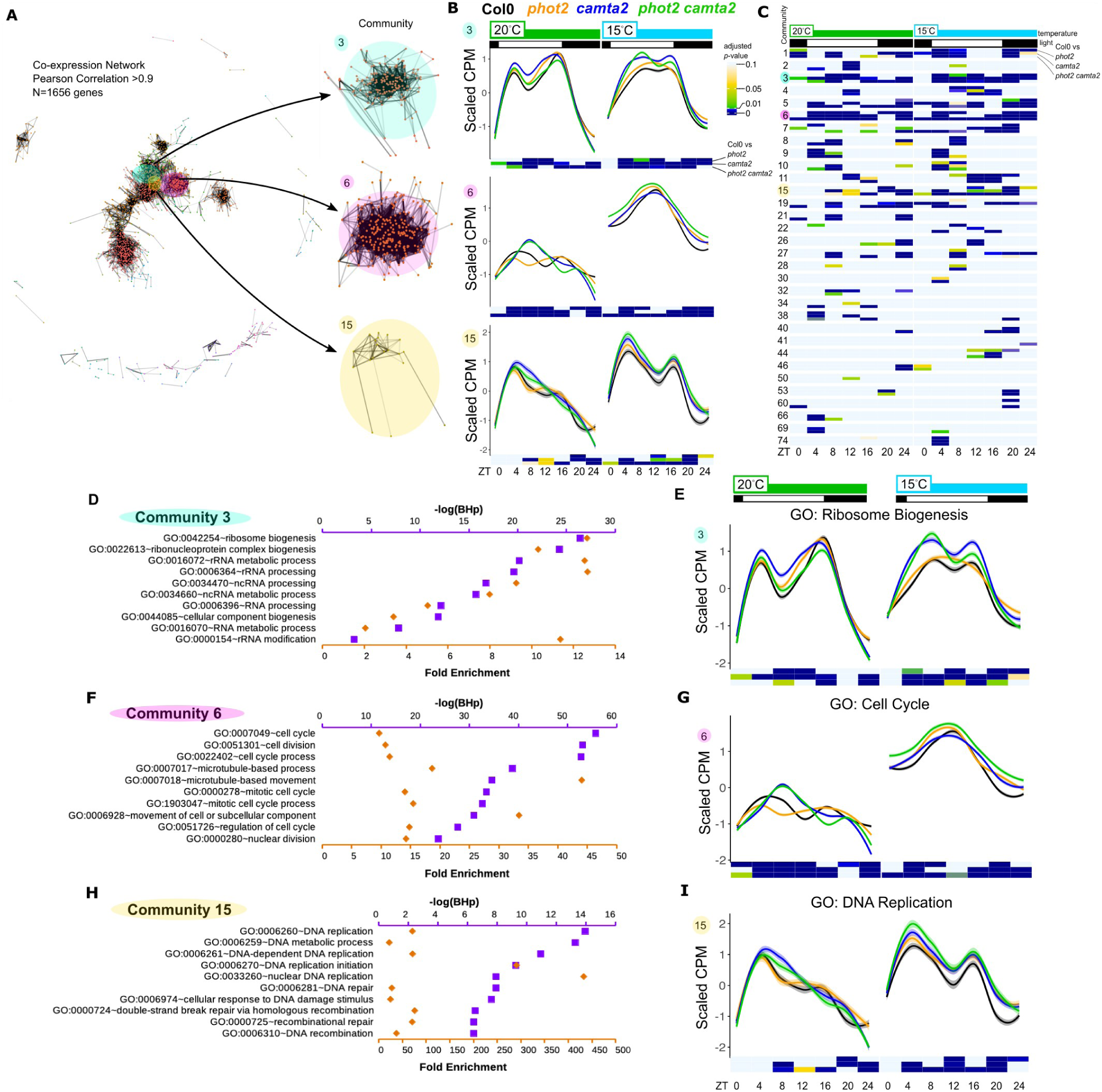
System-level control of basic growth processes by PHOT2 and CAMTA2. (A) Coexpression network constructed from genes with Pearson Correlation Coefficient >0.9 (N=1656 genes). Communities 3, 6, and 15 of the 81 identified communities are magnified and shown at right. (B) Mean scaled timecourse expression patterns for communities 3, 6, and 15 are shown by time of day, genotype and temperature. Traces are colored by genotype. Col-0: black, *phot2*: orange, *camta2*: blue, *phot2 camta2*: green. Shaded area around traces are 95% confidence interval. Black and white bars at top indicate the light-dark cycle.Temperature is shown in green and light blue bars at top. Heatmaps below each graph are *p*-values derived from 2-way ANOVA+Tukey’s HSD test (genotype X time of day within temperature) for each mutant compared to Col-0. Each heatmap has three rows: first is *phot2* vs Col-0, second is *camta2* vs Col-0, and third is *phot2 camta2* vs Col-0 (labeled at right of Community 3). Color key for heatmap is at left, and is the reference for panels B,C,E,G, and I. (C) Heatmap of p-values derived from 2-way ANOVA+Tukey’s HSD test for each mutant compared to Col-0 for all identified co-expression communities made up of at least four genes. Heatmap is divided by temperature (green and blue bars at top) and by community (numbers at left). Color bar at left shows color mapped to p-value. Communities 3, 6 and 15 are highlighted for comparison to (C). (D, F, H) Gene ontology for communities 3, 6, and 15, respectively. The top ten most significant GO-Biological Process terms for each community are shown, with -log(Benjamini-Hochberg p- value) [-log(BHp)] mapped to the upper axis in purple, and Fold Enrichment mapped to the lower axis in orange. (E, G, I) The means of all samples within the top GO Biological Process category from each community are plotted as in (B). (E) Community 3, Ribosome Biogenesis. (G) Community 6, Cell Cycle. (I) Community 15, DNA Replication.

Light input to the PHOT/CAMTA/NPH3 system most likely comes directly through PHOT photosensors, but temperature input is known to effect many parallel processes. Given the links that we have seen between PHOT2 and CAMTA2, and the known functions of CAMTA genes in low temperature-dependent gene expression, we searched our RNAseq data set for temperature-dependent gene expression patterns that were disrupted in the *camta2* mutant. We found that expression of the calcium-dependent lipid binding protein *<u>E</u>N<u>H</u>ANCED <u>B</u>ENDING* 1 (*EHB1*) was up-regulated in lower temperature in Col-0, with reduced peak expression in *camta2* [Fig. 5A]. Expression in *phot2* was similar to Col-0 control under both temperatures. Quantitative RT-PCR in an independent RNA null allele of *CAMTA2* shows that low temperature-dependent induction of *EHB1* transcript is largely dependent on CAMTA2 [Fig. 5B]. We found that an N-terminal fragment of CAMTA2 binds to a 25bp fragment of the proximal EHB1 promoter in-vitro. Both wild-type and mutant (CGCG>ATAT) competitor DNA can partially reduce binding in 200x molar excess, but is unable to bind a mutated probe [Fig. 5C]. This suggests that the core four-base motif is not the only feature recognized by CAMTA2, or a CAMTA2-containing complex. EHB1 acts as a negative regulator of NPH3-dependent phototropic and gravitropic growth via direct interaction with the BTB domain^40^. This, combined with our expression data in the *camta2* mutant, led us to hypothesize that plants lacking EHB1 would show increased TLN under low temperature. We tested two alleles of *camta2* and two alleles of *ehb1* under 25°C and 15°C. These mutants were similar to Col-0 controls under 25°C, but show comparable increases in TLN under 15°C [Fig. 5D]. Together, these data are consistent with a role for EHB1 in transducing temperature information from CAMTA2 to the PHOT pathway, where it modulates NPH3 activity [Fig. 6].

**FIGURE 5:**
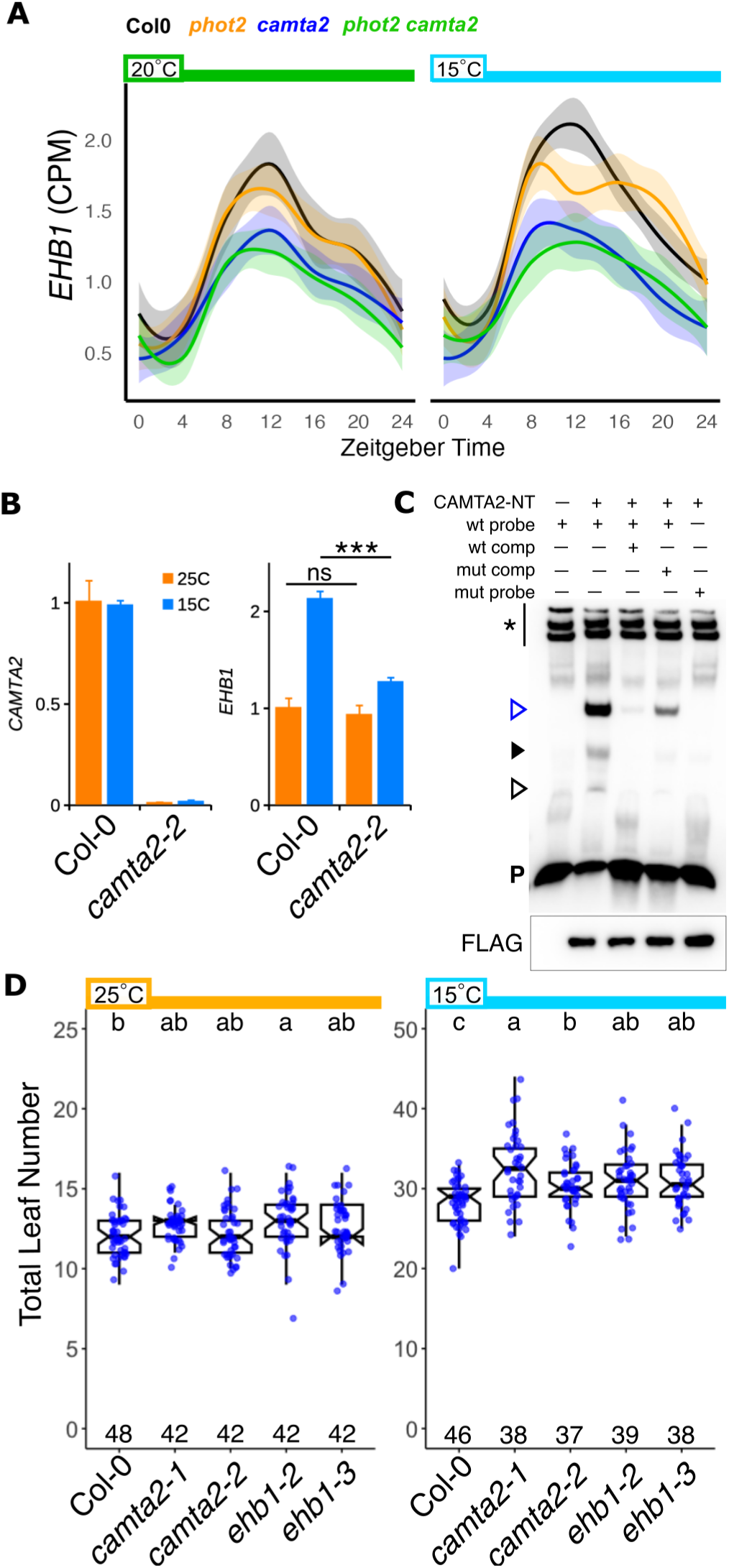
EHB1 connects CAMTA2 temperature inputs to PHOT signaling. (A) EHB1 transcript time courses plotted as Counts Per Million reads (CPM) for plants grown at 20C (green bar) and 15C (light blue bar). Traces are colored by genotype. Col-0: black, *phot2*: orange, *camta2*: blue, *phot2 camta2*: green. Shaded areas around traces are 95% confidence interval. (B) Taqman probe-based qRT-PCR of CAMTA2 and EHB1 in Col-0 and camta2[SALK_139582] (*camta2-2*) under 25C (orange bars) and 15C (blue bars). ns= not significant and *** = p<0.001 by Student’s T-Test. (C) CAMTA2-NT (N-terminus - aa1-364) was expressed in wheat germ extract for Electrophoretic Mobility Shift Assay (EMSA). A 25bp fragment of the EHB1 promoter containing the core CAMTA-binding motif (CGCG) was used as the target sequence. Mutant competitor and probe changed CGCG to ATAT. Unlabeled competitors were supplied at 200x molar excess over labeled probe. The first lane contains unprogrammed wheat germ extract (WGE). *= endogenous biotinylated proteins in WGE; Open blue arrowhead = likely primary complex containing CAMTA2; Solid black arrowhead = light band is non-specific from WGE, with overlapping possible CAMTA2 and wt promoter-dependent complex; Open black arrowhead = possible CAMTA2 monomeric binding event. P = free probe. Done twice with similar results. Lower panel is CAMTA2-NT-FLAG protein in the EMSA reactions. (D) Flowering time assay of *camta2* and *ehb1* mutants. Flowering was assayed as in Fig. 2A. Letters indicate statistical groupings by one-way ANOVA between genotypes, within temperature condition.

**FIGURE 6:**
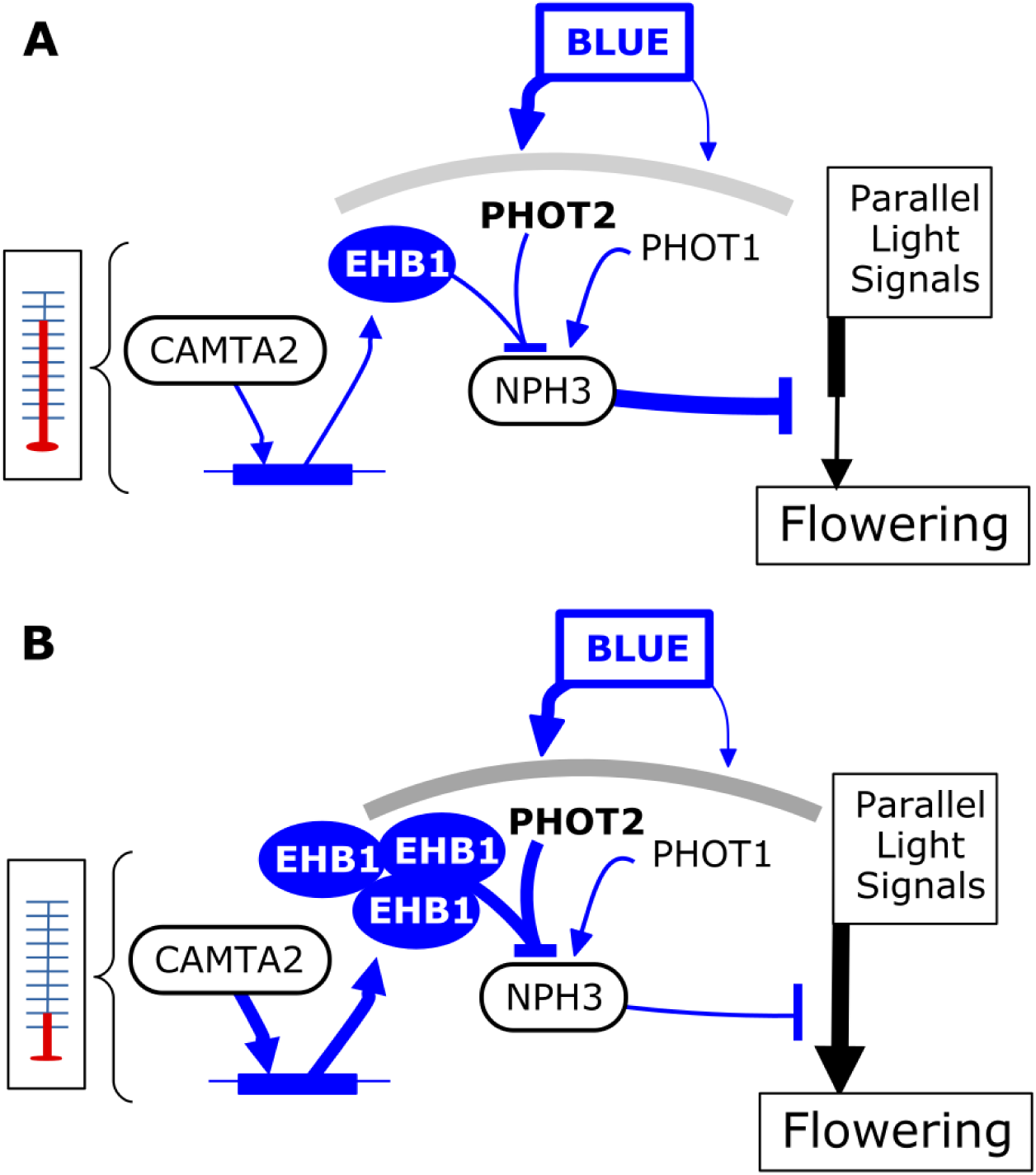
Model of the PHOT2-CAMTA2 light-temperature coincidence detector. A model of the PHOT-CAMTA2-NPH3 light-temperature coincidence detector under warm (A) and cool (B) temperatures. PHOT1 promotes and PHOT2 inhibits the active form of NPH3. NPH3 negatively regulates light signals that promote flowering. CAMTA2-dependent EHB1 accumulation in low temperature inhibits NPH3’s ability to oppose pro-flowering signals. In *phot2* mutants, the active form of NPH3 dominates, strengthening its ability to repress pro-flowering signals. In *camta2* or *ehb1* mutants, EHB1 fails to accumulate, allowing NPH3 to more strongly inhibit pro-flowering signals, thus increasing TLN.

## DISCUSSION

Understanding how plants set and regulate temperature responses to influence developmentally and agronomically relevant traits is increasingly important in a changing climate. In this study, we demonstrate light and temperature dependent control of flowering time by the PHOT2 blue photoreceptor. PHOTOTROPINS have been described as having no role in flowering since their identification^8,18^. Typical laboratory growth temperatures for *Arabidopsis* lie in the 20-23°C range. We observed increased TLN in *phot2* mutants at 15°C, a condition under which *Arabidopsis* grows quite slow and is rarely tested. This may explain why a role for PHOT2 in flowering regulation was overlooked. Our modifications of environmental light conditions under warm and cool temperatures demonstrate PHOT2- and NPH3-dependent light-temperature signal integration. Given that NPH3 gates the influence of PHOT2 [Fig. 2D-F], and the *nph3* early flowering phenotype is light intensity-dependent [Extended Data 2C], we hypothesize that NPH3 acts to inhibit other light-dependent signals that act to promote flowering, and that PHOT2 negatively regulates this function of NPH3 [Fig. 5E].

Thermal and spectral sensitivity of flowering is widely distributed among light signaling pathways. Blue light acts through seven of the 13 non-photosynthetic photoreceptor proteins in *Arabidopsis.* FLAVIN-BINDING KELCH REPEAT F-BOX 1 (FKF1) controls flowering by degrading a transcriptional repressor of CONSTANS (CO)^41–43^. ZEITLUPE (ZTL) and LOV KELCH PROTEIN 2 (LKP2) regulate flowering under short days^44^. Recent work has described blue light via CRYPTOCHROME 2 (CRY2) feeding into temperature-regulated flowering by controlling splicing of flowering regulators in a day-length-dependent manner^45^. Plants lacking red/far-red sensing PHYTOCHROME (PHY) B and/or D proteins flower with fewer leaves than controls in short days under 22°C, but are similar to controls under 16°C^46,47^. PHYB also feeds into a shade-high temperature growth regulation pathway, suggesting multiple light-temperature coincidence detectors operate across different processes^48^. Our data suggest that the PHOT2-NPH3 system may interact with one or more of these photoreceptor signaling pathways, cross-regulating incoming light and temperature information, and/or buffering against fluctuations. Given that reducing the blue component of white light phenocopies delays flowering in Col-0 [Fig2A-C], but increasing blue or red failed to modify the phenotype, [Extended Data 2A-B], we prefer the hypothesis that PHOT2-NPH3 signaling acts to gate the influence of low-intensity blue light signaling pathways. Consistent with this, recent work has indicated that *Arabidopsis* senses gradual light intensity changes at dawn and dusk^49^. This work therefore provides a unique position from which to map out the processing and integration of light and temperature information between photoreception systems.

We characterized flowering time in a suite of mutants covering PHOT-interacting genes, finding a role for NPH3. We show that NPH3 is epistatic to PHOT2 in flowering regulation[Figs. 1A, 2D-F Extended Data 2C]. The NRL family has 33 members in *Arabidopsis*, only a few of which have been characterized experimentally^22^. Upon blue light activation, PHOT1 both directly phosphorylates and triggers dephosphorylation of NPH3, releasing it from the plasma membrane into cytoplasmic complexes^50–54^. The precise function of NPH3 in these cytoplasmic complexes is unknown. PHOT2 and RPT2 subsequently facilitate NPH3’s return to the plasma membrane via unknown mechanisms^52,55^. NPH3 and NCH1 form complexes with CULLIN3 to ubiquitinate target proteins, but whether RPT2 forms similar complexes is unclear^26,56^. Few target or client proteins for NRL proteins have been described, and identification of these interactors will be of distinct interest for understanding cross-regulation of signaling pathways by NRL proteins. RPT2 and NCH1 have redundant functions in regulation of chloroplast movement via JAC1^57,58^. We observe no notable flowering phenotype in plants lacking JAC1 or CHUP1, both of which are critical for chloroplast movement [Fig. 1A]. This indicates that chloroplast movement is highly unlikely to be a central factor in the flowering phenotype.

Multiple links support the hypothesis of signaling between PHOTs through NRL proteins, calcium, and CAMTA TFs. CAMTAs are regulated by intracellular Ca^2+^ and CALMODULIN (CAM) binding^59^. Cold acclimation, in which pre-treatment with low temperature increases survival when exposed to freezing temperature, requires Ca^2+^ mobilization, with application of Ca^2+^ channel blockers or Ca^2+^ chelators resulting in reduced survival in low temperature^60^. The *camta1 camta3* double mutant also shows reduced cold acclimation^24^. A recent study reported NPH3 co-purifying with CAM7, although this interaction was not further examined^50^. A link between PHOT signaling and the CAMTA transcription factors was proposed when NCH1 was identified as a CAMTA3 interacting protein, and the *nch1* mutant was shown to modify CAMTA3-dependent responses to bacterial infection, blocking CAMTA3 ubiquitiniation and degradation^26^. While a subsequent study in protoplasts failed to corroborate this finding, it is possible that this interaction is dependent on cell-specific conditions or factors^30^. However, studies in potato characterized an NRL protein as a susceptibility factor for *Phytophthora infestans* infection, and showed that blue light and PHOT1 decreased the immune response^61,62^. Our findings are in line with the emerging literature suggesting a series of pathways linking PHOTs, NRL proteins, Ca^2+^, and CAMTA transcription factors to regulate development and immunity.

We identify EHB1, a Ca^2+^-dependent lipid binding protein, as a temperature-dependent link between CAMTA2 and the PHOT-NPH3 blue light signaling pathway. Low-temperature-dependent *EHB1* transcription in the mid-day depends on CAMTA2, which can bind directly to the *EHB1* promoter [Fig. 5A-C]. EHB1 was described as a negative regulator of NPH3, likely via direct interaction with NPH3’s BTB domain^40^. EHB1 membrane association is enhanced by calcium^63^. Blue light-activated Ca^2+^ transients rely heavily on PHOTs, with PHOT2 likely to be the dominant activator^64–66^. PHOT2-dependent calcium currents may therefore act on EHB1 to promote NPH3 translocation from cytosol to plasma membrane. The observation that *phot2*, *camta2* and *phot2 camta2* have identical temperature-responsive flowering phenotypes suggests that both factors are necessary for this control mechanism [Figs. 3B, Extended Data 4]. Plants lacking EHB1 or CAMTA2 have identical low-temperature-dependent flowering phenotypes as well, consistent with the hypothesis that EHB1 is the primary factor linking CAMTA2 to PHOT2 [Fig. 5D]. CAMTA TFs are also regulated by calcium through CaM binding to C-terminal domains. It is possible that PHOT2-dependent Ca^2+^ currents can simultaneously enhance EHB1 transcription via CaM-CAMTA2 and membrane localization via Ca^2+^ and EHB1 effects on NPH3, perhaps through membrane binding. The dynamic relationships among PHOT2, calcium, CAMTA2, and EHB1 will be of definite interest for future study.

Studies of PHOT-dependent gene expression have thus far been restricted to examining the effects of acute blue light treatment^67^. The CAMTA family’s effect on transcription is best understood in the context of acute treatment with noxious cold or other stresses^24,35^. Our time-course transcriptomes allow comparisons across the day, and within a relatively narrow ambient temperature band (20°C vs 15°C), rather than noxious high or low temperature. There are six CAMTA TFs in *Arabidopsis*, and CAMTA genes are known to be partially redundant^29,38^. We observe more DEGs in the double mutant than in either single mutant. It is possible that PHOT2 works with more than one CAMTA TF, and that removing CAMTA2 uncovers more extensive regulation through other family members that is further compromised by removal of PHOT2. Our network-based analysis reveals system-level PHOT2- and CAMTA2-dependent regulation of translation, the cell cycle, and DNA replication, among other processes [Figs. 4C-G]. The *phot2* mutant exhibits not only increased TLN, but an increase in leaves generated per day until flowering, consistent with increased organ formation, cell division, and growth [Figs. 2C]. Three types of growth are represented here: translation which increases cytoplasmic volume and cell size, the cell division increases cell number, and DNA replication contributes to both cell division and endoreduplication^68,69^. At whole-shoot resolution, we cannot say how these types of growth are distributed through the plant, and how they are altered in the mutant lines. It will be of considerable interest to examine division rate, DNA content, and cell size in growing tissues in *phot2* and *camta2* mutants.

Systematic changes in environmental light and temperature across the year provide information that plants decode to fine-tune life history traits. Here we describe a role for PHOT2, CAMTA2, and NPH3 in the coordinated processing of blue light and low temperature information, feeding into flowering time control. This coincidence detector is positioned as a negative regulator of other flowering-promoting signals in low temperature, providing new insight into enviromental signal integration. Examination of this system in a variety of plant species will be of interest, as it may provide an access point for engineering tolerance to a range of temperatures and understanding mechanisms of resilience in the face of broad climate shifts.

## MATERIALS AND METHODS

### Plant Material

*Arabidopsis* plants are in the Col-0 background unless otherwise noted. *gl1*, *phot1-5 gl1*, and *phot2-1 gl1* lines were described. *phot1* (SALK_146058), *phot2* (SALK_142275), *nph3* (SALK_110039), *chup1* (SALK_129128C), *jac1* (WISCDsLOX457-490P9), *elf3* (SALKseq_043678.1), *camta1* (SALK_008187), *camta2* (1:SALK_007027, 2:SALK_139582), *camta3* (SALK_001152), *ehb1* (1:SAIL_385_C07, 2:SAIL_1307_C10.v1) mutants were obtained from the Arabidopsis Biological Resource Center (Ohio State University). Mutant combinations were generated by standard genetic crosses and confirmed by PCR genotyping using primers in Supplemental Table 1.

### Flowering Assays

Seeds were surface sterilized using the chlorine gas method in a chemical fume hood for ∼1 hour, then plated on 1/2x LS + 0.8% phytoagar medium (Caisson) and stratified in the dark at 4C for 6-7 days. Seeds were germinated in a growth room set on a long-day (LD16:8) photocycle (light intensity ∼80μmol/m^2^s) at ∼22°C. Seedlings were transplanted to soil with fertilizer and fungal inhibitor on day three (two days after germination) and moved to growth chambers (Percival) under the indicated growth conditions. Light for flowering assays was provided by cool white fluorescent bulbs (Philips) set at the indicated intensities. Light intensity and spectra were confirmed using LI-250A light meter or LI-180 spectrometer (LiCor). Supplemental blue and red light were provided by LED bulbs in addition to the fluorescents. Blue light was specifically reduced by filtering the white fluorescent light with #101 Yellow filter sheeting (Lee Filters). Chamber ambient temperatures were confirmed with a glycerol thermometer (Fisher Scientific) or HOBO temperature loggers. Trays were rotated and cycled around the chamber every 1-2 days to avoid position effects. Flowering time was scored at the appearance of buds at the center of the rosette. Leaves were counted under a dissecting microscope (Leica).

### RNA isolation and RNA sequencing

RNAseq1: Plants were sterilized, stratified, germinated, and transplanted as for the flowering assays. Above-ground tissue from three seedlings per sample were collected and snap-frozen in liquid nitrogen in biological duplicate at ZT16 and ZT24 of day 14 post-germination. Total RNA was extracted with the Qiagen RNeasy Plant mini kit according to manufacturer’s instructions. Single-end sequencing libraries were made and sequenced on the HiSeq2500 at the Salk Institute Next Generation Sequencing core.

RNAseq2: Plants were sterilized, stratified, germinated, and transplanted as for the flowering assays. Above-ground tissue from six seedlings per sample were collected and snap-frozen in liquid nitrogen in biological triplicate at ZT0, 4, 8, 12, 16, 20,and 24 of day 10 post-germination. Total RNA was extracted with the RNeasy Plant mini kit (Qiagen) and DNAse treated (Qiagen) according to manufacturer’s instructions. RNA was poly-A selected and paired-end libraries were prepared at the Salk Institute Next Generation Sequencing facility. Sequencing data were generated on a NovaSeq 6000 S4-XP with PE100 reads at the University of California, San Diego sequencing core. Samples were demultiplexed using bcl2fastq Conversion Software (Illumina, San Diego, CA).

### RNA-seq analysis

RNAseq1: Reads were aligned to the TAIR10 *Arabidopsis* genome using TopHat, and reads per gene quantified using HTSeq^70^. Differential expression analysis was carried out in edgeR, with comparisons set between mutant and wild type samples within each time point and temperature condition at False Discovery Rate (FDR) =0.05 and Fold Change (FC)=|0.25|^71^.

RNAseq2: Reads were aligned to the TAIR10 *Arabidopsis* genome using STAR with default parameters except --alignIntronMax 2000 and --outFilterMismatchNmax 2. Reads per gene were calculated using the Araport11 list of genes and transposons in HTSeq with default parameters except “-m intersection-strict -s reverse -r pos”. Differential expression analysis between mutant and wild-type samples at each time point was carried out in edgeR with False Discovery Rate (FDR) ≤0.05 and Fold Change (FC)=|0.25|. Genes with cumulative CPM>1 across all samples were retained for analysis.

### Network co-expression analysis

Network co-expression analysis was carried out in R as described, starting from the complete list of all genes identified to be differentially expressed in any of the mutant lines at any time-point relative to Col-0 (N=3478 genes - see Supplementary Data 3^72^. To more directly compare gene expression dynamics across the day, CPM values were scaled such that the mean expression across all samples for each gene is equal to 0. Pearson Correlation Coefficients (PCC) were calculated between each gene across all samples using the “cor” function (R). Genes that were co-expressed with PCC>0.9 were used to construct a network using the igraph R package^73^. Co-expressed communities within this network were identified using “cluster_edge_betweenness” in igraph. Gene Ontology enrichment analysis by community membership was carried out in DAVID (https://david.ncifcrf.gov/home.jsp) with an inclusion threshold of Benjamini-Hochberg *p*-value<0.05^74,75^.

### qRT-PCR

RNA was extracted from six 10-day-old soil-grown seedlings with the RNeasy Mini kit (Qiagen). First strand synthesis was carried out with 2ug total RNA using the Maxima cDNA kit according to manufacturer’s instructions. Real-Time PCR was done using TaqMan Gene Expression Assays (Thermo - Supplementary Table 1) and TaqMan Gene Expression Master Mix according to manufacturer’s instructions. Reactions were carried out in a BioRad CFX Opus 384 Real-Time PCR Machine. Relative transcript abundance was determined by the ΔΔCq method using the static AT4G26410 transcript as the internal control.

### Electrophoretic Mobility Shift Assays

A plasmid encoding CAMTA2-NT (amino acids 1-364) driven by the SP6 promoter was synthesized (GenScript) and 4μg of plasmid expressed in the TnT Wheat Germ Extract Kit (Thermo) for 2.5h at 25°C. Two μL of programmed or unprogrammed extract was used per reaction. A 23bp fragment of the EHB1 promoter—biotin labeled probe and unlabeled competitor—were synthesized (IDT). Binding was carried out using the LightShift Chemiluminescent EMSA Kit (Thermo) according to manufacturer instructions, with the addition of 2.5% glycerol, 5mM MgCl_2_, and 3μg double-stranded polyAT DNA, with 20fmol labeled probe and 4pmol (200x) unlabeled competitor per reaction as indicated. Detection was carried out with the Chemiluminescent Nucleic Acid Detection Module (Thermo) according to manufacturer instructions and imaged on a Sapphire Biomolecular Imager (Azure Biosystems).

### Western Blots

Samples from the EMSA reactions were mixed with 2xLDS buffer (LIfe Technologies) and heated for 5 min at 85C. Samples were run on pre-cast 4-12% gel (Life Technologies) and transferred to nitrocellulose membrane. Membranes were blocked with TBST (tris-buffered saline + 0.2% Tween-20) with 3% milk, and incubated with monoclonal anti-FLAG-HRP M2 (Sigma, #A8592) at 1:2000 dilution in blocking solution for 1h shaking at room temperature. membrane was washed 5x in TBST, then 2x in water. Detection was carried out with SuperSignal West Pico PLUS (Pierce) according to manufacturer instructions. Membrane was imaged on a Sapphire Biomolecular Imager (Azure Biosystems).

### Statistics

Sample numbers are indicated within each figure. Comparisons between two groups were done with t-tests (two sided), and comparisons among three or more groups were done with one-way ANOVA and Tukey’s HSD test at alpha=0.05. Genome X Environment interactions were tested by two-way ANOVA and Tukey’s HSD test at alpha=0.05. Leaf Number Ratio (LNR) was calculated as described^34^. Significant overlap between populations, including binding site enrichment and comparison of DEG lists between genotypes, was determined using Fisher’s Exact Test.

## AUTHOR CONTRIBUTIONS

A.S. and J.C. conceptualized the study, designed experiments and wrote the paper, A.S. performed all experiments, and analyzed the data. J.C. acquired funding and supervised.

## ACKNOWLEDGEMENTS

We thank Dr. John Christie, Dr. Claudia Oecking, and Dr. Michael Thomashow for sharing information and reagents prior to publication. J.C. is an investigator for the Howard Hughes Medical Institute. This work was supported by National Institutes of Health (NIH) grant R35-GM122604 to J.C., and by the Howard Hughes Medical Institute. A.S. received support from the Salk Institute Pioneer Postdoctoral Endowment Fund. We thank Andrew Gregory, Megan Rae and Ashnah Shwany for technical assistance. This work was supported by the Next Generation Sequencing Core Facility at the Salk Institute with funding from NIH-National Cancer Institute Cancer Center Support Grant P30 014195, the Chapman Foundation and the Helmsley Charitable Trust. This publication includes data generated at the University of California, San Diego Institute of Genetic Medicine Genomics Center utilizing an Illumina NovaSeq 6000 that was purchased with funding from NIH Shared Instrumentation Grant S10 OD026929.

## EXTENDED DATA LEGENDS

**Extended Data 1:**
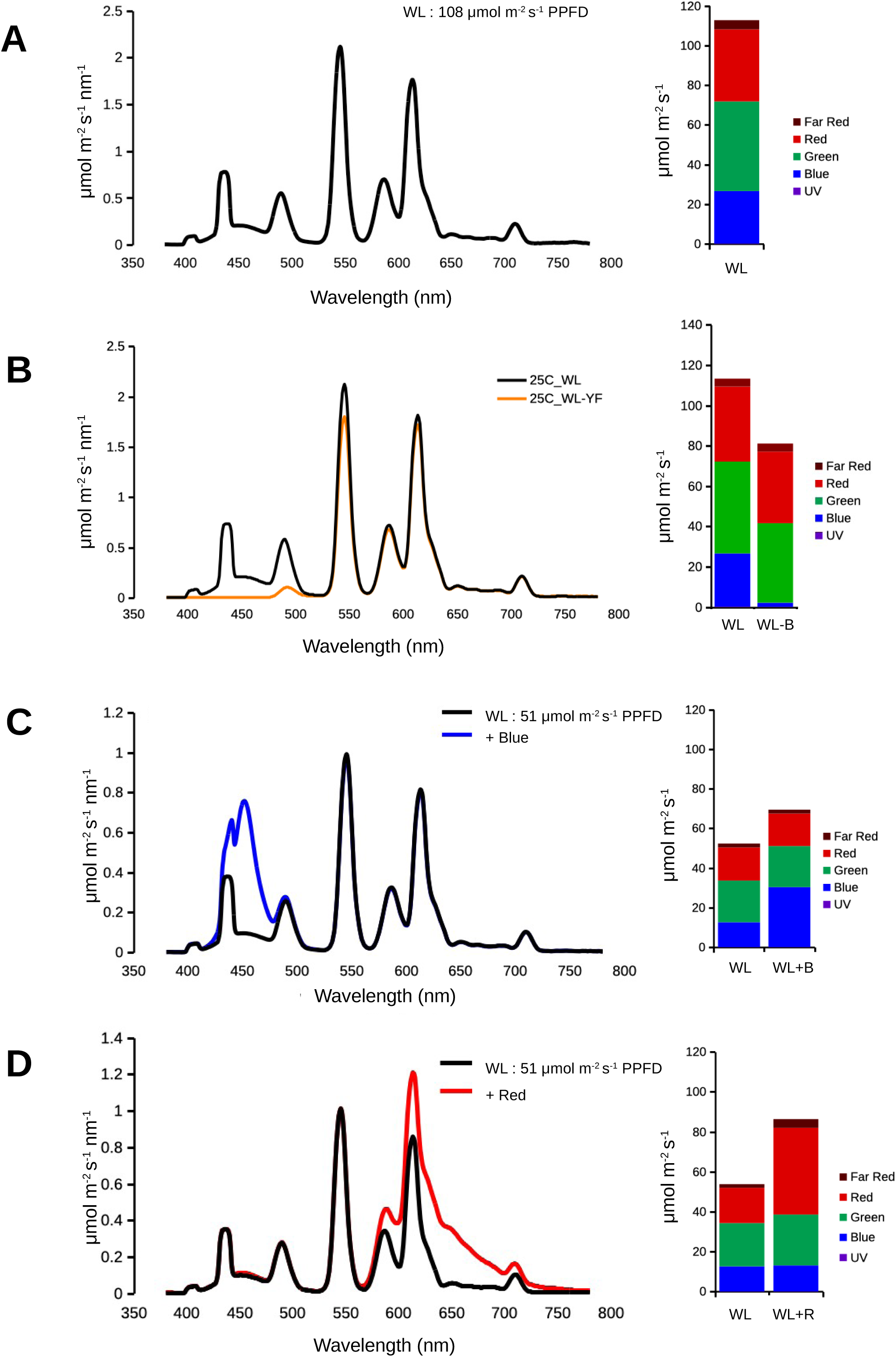
Representative light spectra for flowering and RNAseq experiments. Bar charts to the right of each spectrum trace shows a breakdown of UV, Blue, Green, Red and Far Red components. (A) Representative of the light conditions used for most flowering experiments, unless noted, and both RNA sequencing experiments. (B) White light 105 μmol m^2^ s^-1^ PPFD vs yellow filtered light. (C) White light 51 μmol m^2^ s^-1^ PPFD +/- blue LED supplement. (D) White light 51 μmol m^2^ s^-1^ PPFD +/- red LED supplement.

**Extended Data 2:**
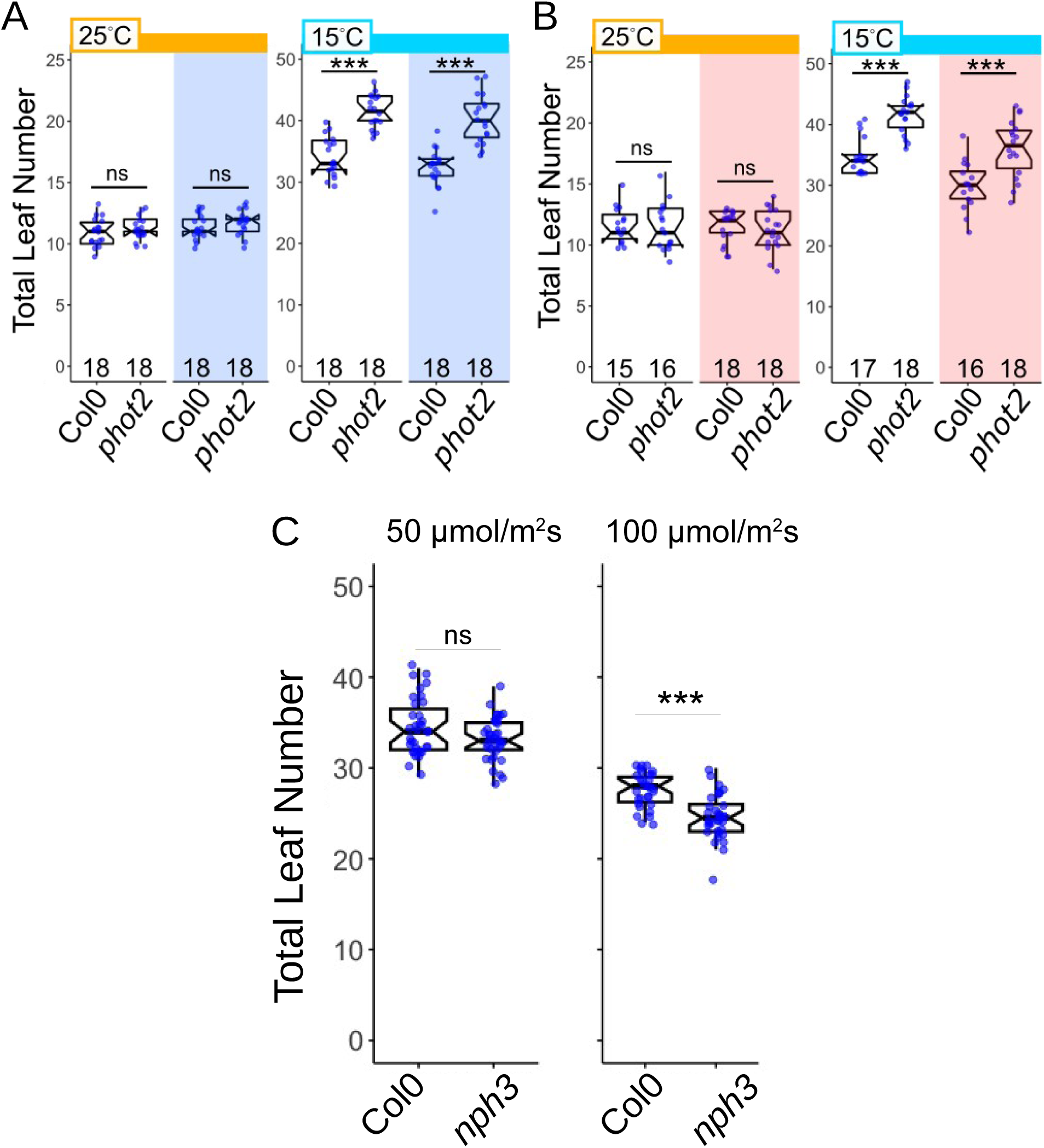
Genetic and Environmental conditions modify TLN at flowering in *phot2* and *nph3*. (A-B) Excess Blue (A) or Red (B) light does not modify the TLN phenotype of *phot2*. Flowering time as assayed by total leaf number at the time of bud appearance in Col-0 and *phot2* under the indicated light and temperature conditions. Colored bars above boxplots indicate temperature: orange = 25C, light blue = 15C. Light conditions are indicated by background shading: white = white fluorescent light ∼50 m-2 s-1, blue = white fluorescent + blue LED, red = white fluorescent + red LED. See Extended Data 1 for spectra. White light data from Col0 and *phot2* in (A) and (B) were combined and plotted in Figure 2D, and further analyzed in Figures 2E and 2F. (D) Light intensity sensitivity in *nph3*. Data for Col-0 and *nph3* are replotted from Figure 2B (50 μmol m2 s-1), and Figure 1A (100 μmol m2 s-1). TLN boxplots are presented as in Fig. 1. Different letters indicate statistically significant difference between genotypes (α=0.05), within each temperature condition by ANOVA+Tukey HSD.

**Extended Data 3:**
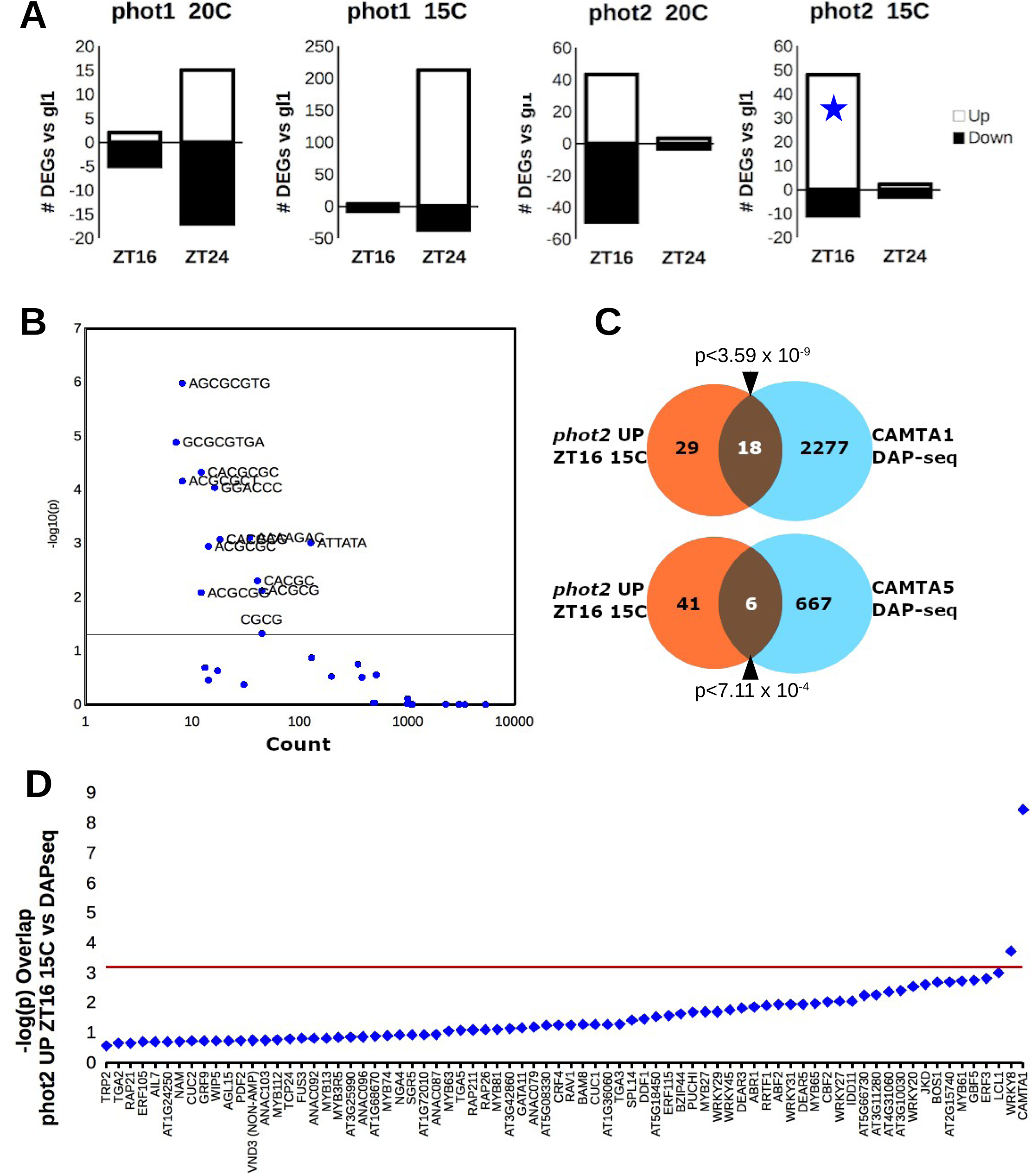
RNA sequencing of *PHOT* mutants suggests a connection between *PHOT2* and *CAMTA* transcription factors. (A) Bar charts show the number of differentially expressed genes (DEGs) identified relative to control (*gl1*) in *phot1-5 gl1* and *phot2-1 gl1* strains grown in soil at either 20°C or 15°C, and sampled at ZT16 (end-of-day) or ZT24 (end-of-night) of day 14 in biological duplicate. Blue star indicates the gene set analyzed further in (B-D). (B) Enrichment of potential cis regulatory elements in 1kb promoter elements of the set of genes up-regulated in *phot2 gl1* in 15°C, ZT16 calculated using the ELEMENT tool. X-axis is the total number of occurrences of the element in the promoters of the gene set. Y axis is the corrected P-value of the enrichment. (C) Genes bound by CAMTA1 or CAMTA5 by DAP-seq and the *phot2* 15°C ZT16 DEG set. P-value of overlap from Fisher’s exact test is shown. (D) Evaluation of DAP-seq target overlap analysis was carried out with 75 other transcription factors in the database with a similar number of reads as CAMTA1 (2000-3750 reads), with each TF represented by a blue diamond and ordered by increasing -log(p) value (Fisher’s exact test). CAMTA1 is the last and most significantly enriched transcription factor on the plot.

**Extended Data 4:**
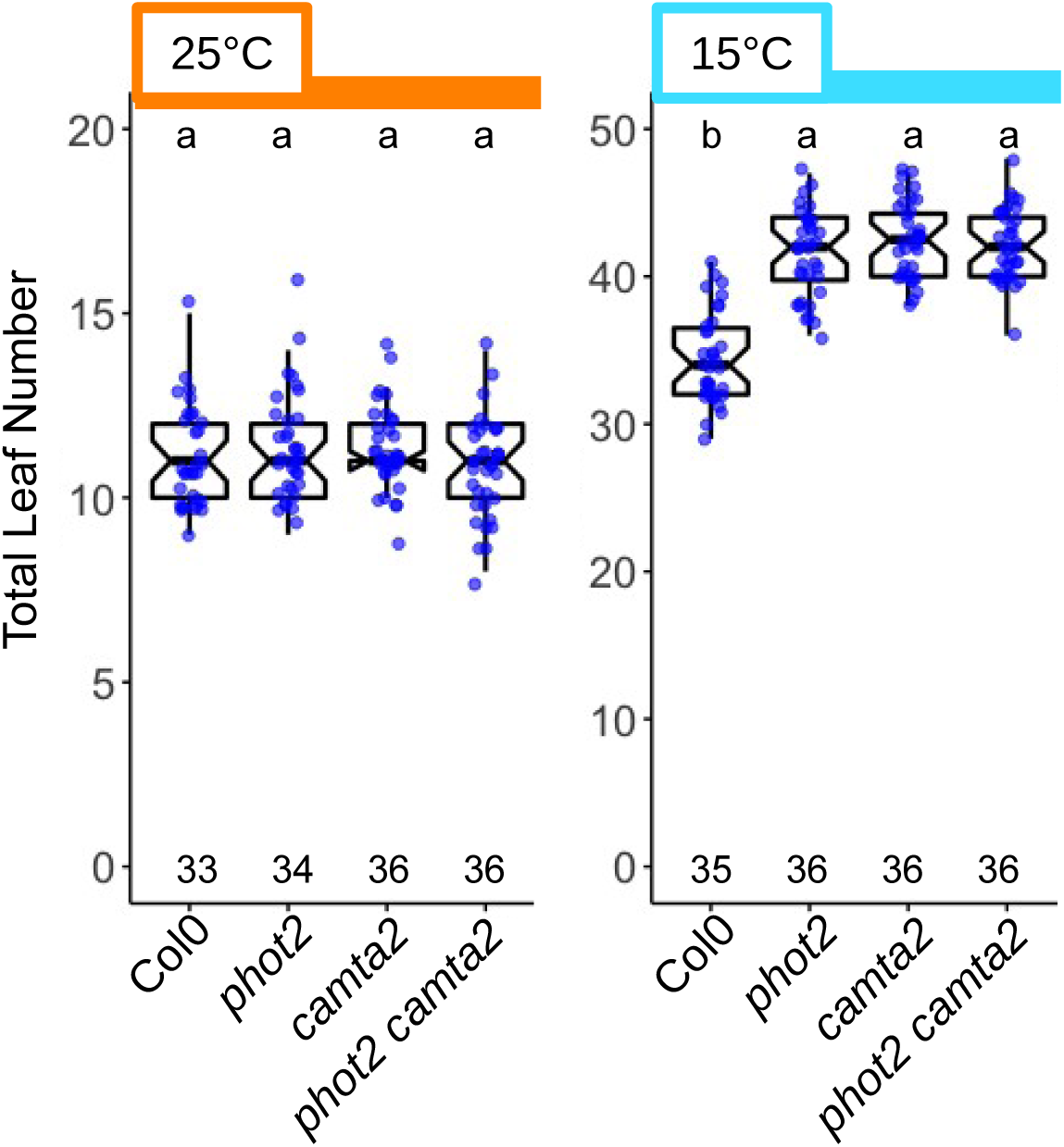
Temperature responsive flowering of *phot2*, *camta2*, and *phot2 camta2* mutants under low light. (A) TLN of Col0, phot2, camta2, and phot2 camta2 plants counted under 50 μmol m-2 s-1 white light at 25°C and 15°C. Col0 and phot2 data are re-plotted from Extended Data Figure 2A-B (pooled white light) for reference.

**Extended Data 5:**
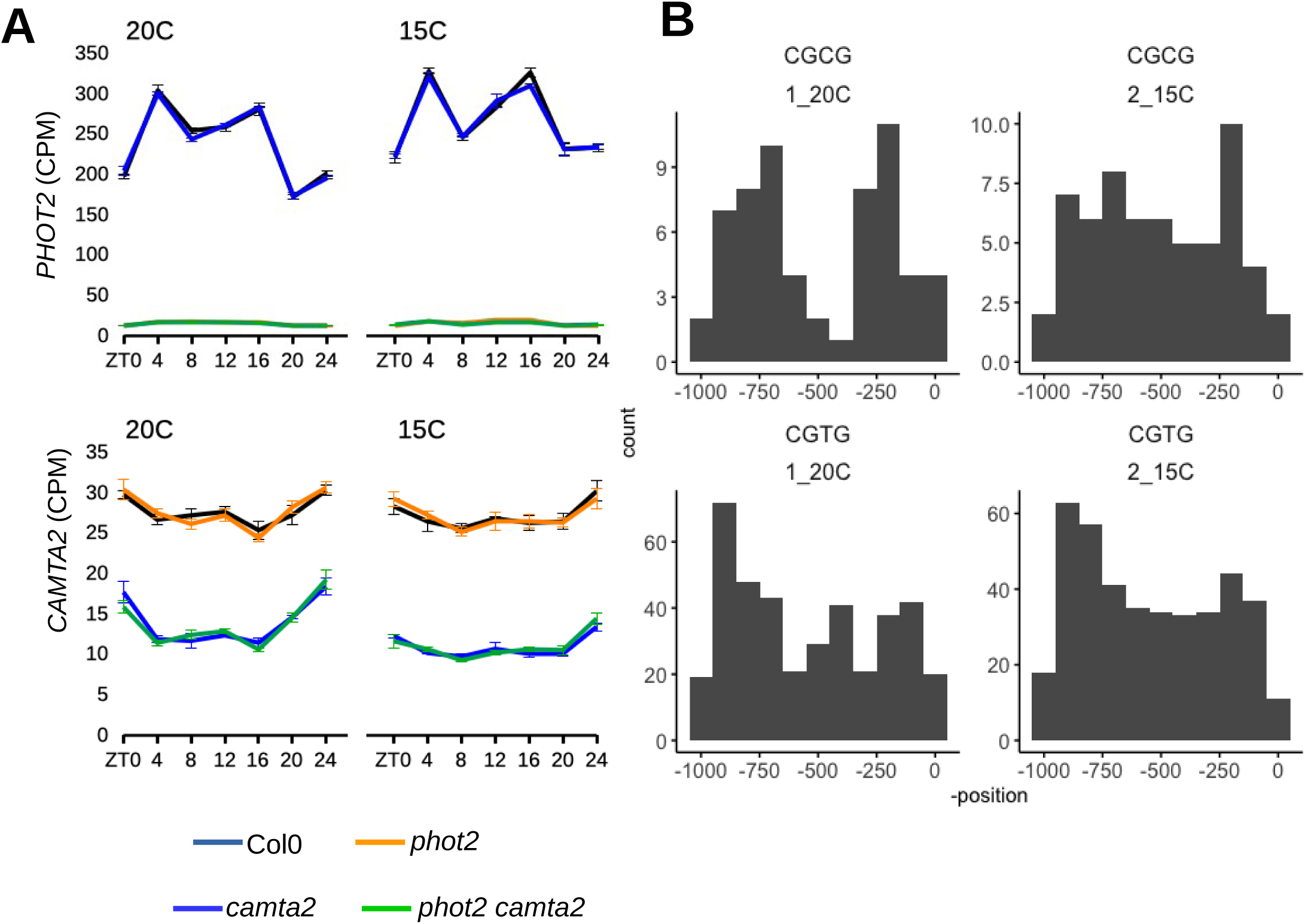
Expression levels and dynamics of PHOT2 and CAMTA2, and likely CAMTA targets. (A) Counts Per Million reads (CPM) for PHOT2 and CAMTA2 transcripts in Col-0, phot2, camta2, and phot2 camta2 plants. Error bars are SEM. (B) Histograms depicting the position of CGCG and CGTG motifs in the promoters of Common DEGs. X axis in in base pairs of the 1kb promoter, with 0 being the transcription start site.

**Extended Data 6:**
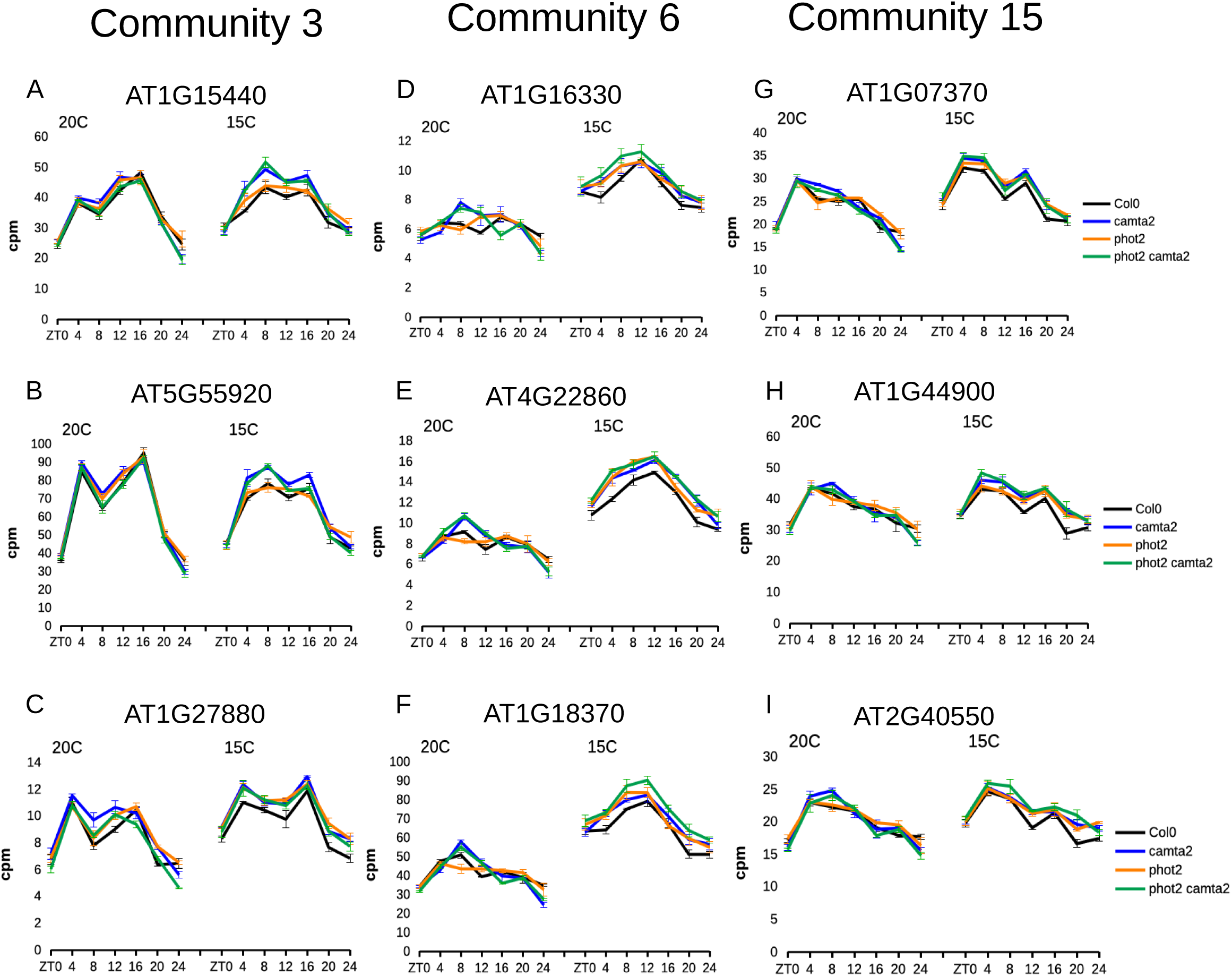
Representative gene expression profiles from communities 3, 6, and 15 derived from network analysis. (A-C) CPM values of genes within the Ribosome Biogenesis GO category. (A) AT1G15440,*PERIODIC TRYPTOPHAN PROTEIN 2* (*PWP2), (B) AT5G55920, OLIGOCELLULA 2* (*OLI2*), (C) AT1G27880, DEAD/DEAH-box RNA helicase. (D-F) CPM values of genes within the Cell Cycle GO category. (D) AT1G16330 CYCLIN B3;1, (E) AT4G22860, TPX-like PROTEIN 3 (*TPXL3*), (F) AT1G18370, NPK1-ACTIVATING KINESIN 1 (NACK1), (G-I) CPM values for genes in the DNA Replication GO category. (G) AT1G07370, *PROLIFERATING CELL NUCLEAR ANTIGEN 1* (*PCNA1*), (H) AT1G44900, *MINICHROMOSOME MAINTENANCE 2* (*MCM2*), (I) AT2G40550, *E2F TARGET GENE 1* (*ETG1*).

**Extended Data 7:**
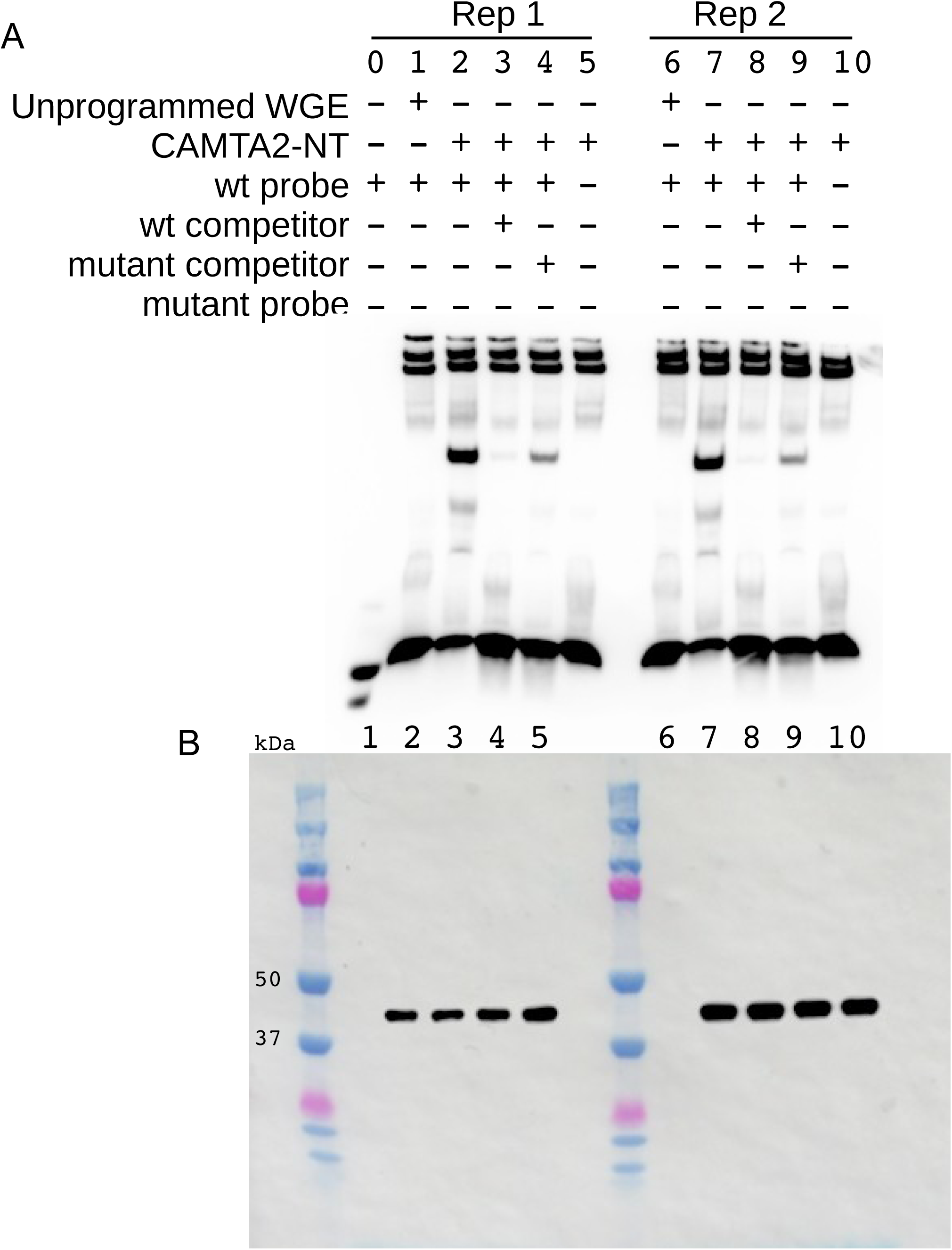
Uncropped gel image from EMSA replicates from Figure 5C. (A) Lane 0 in Rep 1 is only wt probe, no protein or extract, for the purpose of orienting the gel image. Unprogrammed WGE is wheat germ extract without CAMTA2 expression plasmid. Replicates were done using independent CAMTA2 protein preparations. (B) Uncropped anti-FLAG western blot image of CAMTA2-NT-FLAG protein in the EMSA reactions. Shown overlayed with the image of the protein ladder.

